# Epicranial electrical stimulation improves non-navigational spatial memory in macaque monkeys

**DOI:** 10.64898/2026.02.12.705248

**Authors:** Noa Peeleman, Myles Mc Laughlin, Tom Theys, Mathieu Vandenbulcke, Peter Janssen

## Abstract

**Background:** The hippocampus and medial temporal lobe are crucial for spatial memory, and their dysfunction is linked to Alzheimer’s disease (AD), with changes detectable even in preclinical stages. Recently, neuromodulation has gained interest as a potential treatment due to its beneficial effects on AD pathology and cognitive performance. However, outcomes vary significantly based on stimulation parameters and study conditions, and evidence from large animal models remains limited.

**Objective:** To assess whether epicranial current stimulation (ECS) at 40 Hz can improve non-navigational spatial memory and hippocampal activations.

**Methods:** Three rhesus macaques were implanted with spiral platinum electrodes bilaterally on the skull and were trained in a non-navigational spatial memory task. ECS was applied at 40 Hz or at 10 Hz and performance across multiple sessions was evaluated. We further performed ECS during fMRI to examine the spread of activations caused by ECS across the brain in a block-design experiment.

**Results:** ECS at 40 Hz improved performance in a non-navigational spatial memory task, while 10 Hz ECS had minimal or negative effects. Concurrent ECS-fMRI showed extensive brain activations at 40 Hz, including significant hippocampal activations, which was not observed at 10 Hz.

**Conclusions:** Our results show that ECS could be a minimally-invasive and effective approach to improve memory performance and activate the hippocampus. ECS could represent a potential treatment for patients suffering from memory impairment.

## 1 Introduction

The hippocampus and associated medial temporal lobe (MTL) structures are critical for spatial memory [1,2], with their structural and functional integrity essential for learning and memory consolidation [3]. Pathological changes in the hippocampus and MTL are linked to Alzheimer’s disease [4] and can be detected even at preclinical stages [5]. In recent years, neuromodulation has gained interest as an alternative treatment to regulate abnormal brain activity related to the disease progression [6]. While several studies report memory enhancement with brain stimulation [7–10], the outcome is highly dependent on the site of stimulation, behavioral tasks used for evaluation, and the specific stimulation parameters. Not all studies show positive effects, with some studies even reporting memory impairment [11,12].

Recent research highlighted the role of 40 Hz stimulation in the hippocampus [13–16]. Gamma-frequency oscillations (25-100 Hz) have been linked to several higher order cortical processes such as attention, sensory binding and long-term memory [17–19], (see Mann and Paulsen (2005) for detailed review on mechanisms of gamma oscillations in the hippocampus [18], and decreases in gamma oscillations are observed in Alzheimer’s disease before significant amyloid plaque accumulation or cognitive decline [20,21]. Several studies in rodents have demonstrated that 40 Hz optogenetic and sensory light stimulation - but not 80 Hz - can induce gamma entrainment in the hippocampus, promote clearance of amyloid-beta, preserve neuronal and synaptic density, and enhance cognitive performance [14,20], but evidence obtained in a large animal model is currently lacking. We investigated the effects of 40 Hz epicranial current stimulation (ECS) through subcutaneous electrodes implanted on the skull of non-human primates (NHPs) in a non-navigational spatial memory task [22]. By placing the electrodes directly on the skull and isolated from the overlying tissues, we avoided shunting through the skin, achieved higher electric fields in the brain and required a less invasive surgery compared to DBS. ECS at 40 Hz caused significant frequency-specific improvements in spatial memory performance in both animals and induced widespread functional activations in cortical and subcortical structures including the hippocampus, highlighting the potential of ECS as a treatment for memory impairment in patients.

## 2 Methods

All surgical and experimental procedures were performed in accordance with the National Institute of Health’s Guide for the Care and Use of Laboratory Animals and the EU Directive 2010/63/EU, and approved by the Ethical Committee at KU Leuven (project number P075/2021).

### 2.1 Subjects and surgical procedures

Three rhesus monkeys (*Macaca mulatta*; Monkey T: male, 11 kg, 12 years old, Monkey P: male, 9 kg, 10 years old, Monkey C: female, 6 kg, 20 years old) were implanted with a head post (Crist Instruments) fixed to the skull with ceramic screws and dental acrylic. All surgical protocols were performed under propofol anesthesia (10 mg/kg/h) and strict aseptic conditions. After behavioral training, each monkey underwent another surgery in which two custom-built spiral platinum electrodes (6 mm diameter, impedance 0.8-1 kΩ) were implanted bilaterally on the skull, below the zygomatic arch. Electrode location was verified using a post-surgery CT scan (Figure 1A and B).

**Figure 1.**
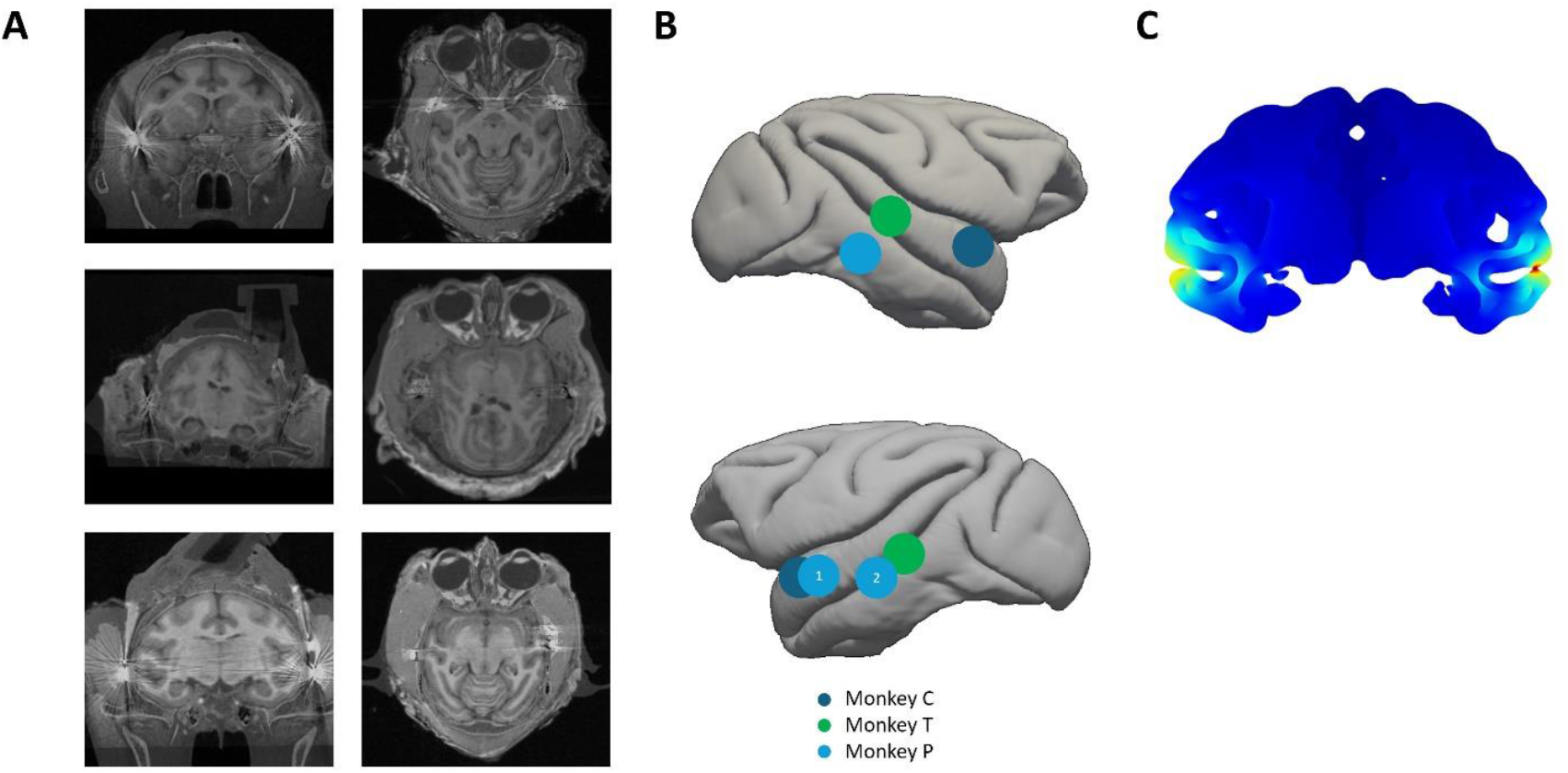
(A) Coregistered coronal (left column) and horizontal (right column) MRI-CT images of monkey C (top row), monkey T (middle row), and monkey P (bottom row) (B) Localization of epicranial electrodes overlayed on an inflated macaque brain. Note that monkey P was implanted twice (indicated as 1 and 2 in the blue circles) (C) Electric field modeling of 3 mA ECS

### 2.2 Behavioral training

Two monkeys (T and P) were trained in a non-navigational spatial memory task, based on the hippocampus-dependent Hamilton Search task [22]. Each session, the monkey sat in a dark room facing a touch screen with its hand on a resting position (Figure 2). The animal had to maintain fixation within a 3×3° window centered on a white dot at the center of the screen. After 500 ms, two targets appeared simultaneously at one of eight possible locations for 500 ms. Following a 2 to 3 seconds delay, the fixation dot turned red (go cue), prompting the animal to touch the previous target locations within 3000 ms. If the animal touched the screen in an area of 4° around the two target locations in any order, it received a liquid reward (Figure 2). In total, there were 28 different combinations of targets (patterns). Patterns included in the study had a performance between 20 and 80% correct to allow for improvement. The resting position of the hand was monitored using an infrared laser beam which was interrupted when the hand was in the resting position. Trials were aborted if the hand moved before the go cue. An infrared-based camera system (EyeLinkII, SR Research) continuously monitored the eye position to ensure the monkey fixated properly.

**Figure 2.**
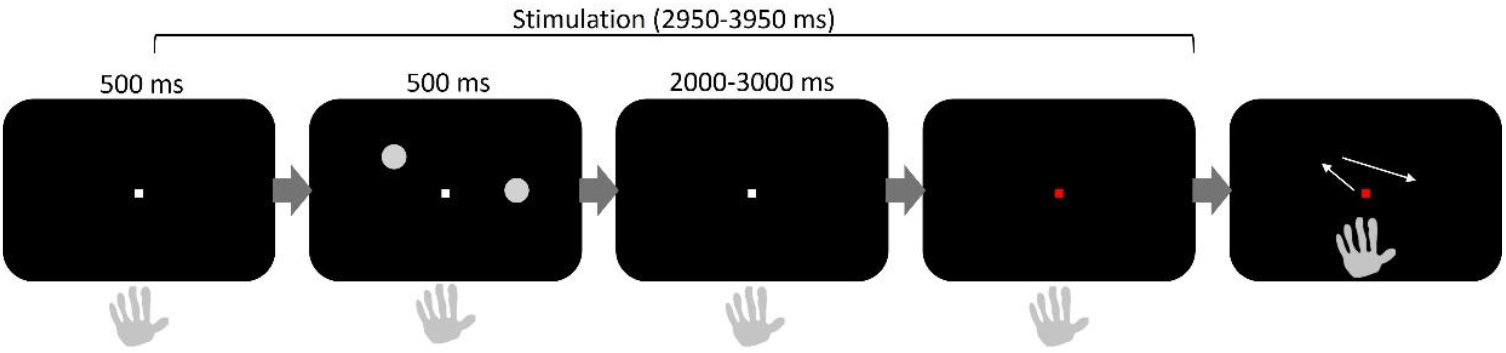
Schematic representation of the temporal sequence of the memory task. The durations of all events (baseline period, stimulus presentation, stimulation duration, delay period) are indicated above the panels. The go-cue was the change in color of the fixation point

After intensive training, once he monkeys’ performance stabilized, we recorded 7 sessions per stimulation condition (40 Hz or 10 Hz) in each animal. Each session consisted of four consecutive blocks, starting with sham stimulation (0.1 mA) and alternating with high-intensity stimulation (3 mA). Blocks consisted of 100 correct trials for monkey T and 70 for monkey P. Ten patterns were selected beforehand and split into two groups, ensuring equal performance between the groups. One group was randomly selected to receive stimulation while the other served as control. The same patterns were used in all 7 consecutive sessions, and the stimulated and nonstimulated patterns were the same in every session. Stimulation started 450 ms before target onset and stopped when the go cue appeared. Stimulation duration ranged from 2950 ms to 3950 ms (on average: 3450 ms), depending on the timing of the go cue. For monkey T, only four patterns were included in the stimulated group in the 10 Hz ECS condition. We also recorded four sessions in monkey P at 80 Hz as an additional control.

### 2.3 Stimulation protocol

Stimulation trains were composed of biphasic square-wave pulses consisting of an anodal leading pulse followed by a cathodal pulse (total 0.48 ms pulse width). Stimulation intensity was set at 3 mA as a previous study conducted in our lab showed that ECS was capable of affecting both spike rate and neural synchronization [23]. For the behavioral study, stimulation was delivered by a function generator (Tektronix AfG3022B; Beaverton, OR USA) in combination with a stimulus isolator (AM systems model 2200; Sequim, WA USA). We modeled the electric field strength induced by 3 mA ECS in the macaque brain using COMSOL (Figure 1C). The electric conductivities of the different model parts were set as follows: CSF (1.65 S/m), GM (0.27 S/m, cerebellum (0.2 S/m), WM (0.127 S/m), skull (0.01 S/m), dental cement (0.01 S/m), and electrode contact (5*10^7^). A detailed explanation of the electric field estimation can be found in Asamoah et al. 2025 [23]. To validate the model experimentally, we measured the pulse artifact for 6 stimulation intensities (0.1 mA, 0.2 mA, 0.3 mA, 0.4 mA, 0.5 mA, and 1 mA) at 5 recording depths and two positions in the lateral-medial direction. The electric field was calculated by dividing the difference in amplitude of the pulse artifact by the distance spacing between two different positions. Stimulation amplitude was adjusted to a stimulation amplitude of 2 mA.

### 2.4 fMRI experiments

Functional images were acquired with a 3 Tesla full-body scanner (Siemens), using a gradient-echo T2*-weighted echo planar imaging sequence of 40 horizontal slices (repetition time [TR], 2 s; echo time [TE], 18 ms; 1.25×1.25×1.25 mm^3^ isotropic voxels) with a custom-built 8-channel phased-array receive coil, and a saddle-shaped, radial transmit-only surface coil. Before each scanning session, the monkeys were sedated with a mixture of ketamine (Nimatek, Eurovet; 12.5 mg/30 min) and medetomidine (Domitor, Orion; 0.25 mg/30 min), and a contrast agent monocrystalline iron oxide nanoparticle (Molday ION, 9-12mg/kg), was injected intravenously [24].

We used a block design alternating between nonstimulation and stimulation blocks (each lasting 40 s), with each run lasting 245 NMR-pulses (490 s). We stimulated at 3 mA and with a frequency of either 40 or 10 Hz, identical to the behavioral experiment. The duration of the stimulation train was fixed at 3 s every 8 s. In all sessions except one, an 8-channel WPI digital stimulator (DS8000, World Precision Instruments; Sarasota, FL USA) was used in combination with a stimulus isolator set on current mode (DLS100, World Precision Instruments). In one session, the same stimulator setup was used as in the behavioral study. The duty cycle per run was 18.37% (90 s of stimulation per run (total run time: 490 s)). Each stimulation train delivered 172.8 μC at 40 Hz and 43.2 μC at 10 Hz. The corresponding charge densities were 28.8 μC/mm^2^ at 40 Hz and 7.2 μC/mm^2^ at 10 Hz.

### 2.5 Data analysis

All data analysis was performed in Matlab R2021a (MathWorks, Natick, MA, USA) using custom-written scripts, unless stated otherwise.

#### 2.5.1 Behavioral analysis

Mean performance was calculated by dividing the total number of correct trials by the total number of correct and incorrect/no response trials, excluding trials with aborted fixation. This was done separately for the group of nonstimulated and stimulated patterns, and performance differences were assessed using Chi square tests of independence. Next, we fitted a generalized linear mixed-effects model (GLMM) with fixed effects for Frequency (40 Hz vs 10 Hz), Stimulation (nonstim vs stim), Block (sham1, high1, sham2, and high2), and Session (1-7) and random intercept for Monkey (T or P). Session was included as a fixed effect to account for day-to-day learning effects. We modeled Block as four separate levels rather than pooling the two sham blocks and the two high stimulation blocks, to quantify the effect of stimulation in each block. Reaction times were calculated as the time needed for the animal to lift its hand after the go cue appeared. Running averages (25-trial window and 10-trial step size) were calculated to visualize the progress in performance across the blocks. Differences in the percentage correct from the running averages were tested using Wilcoxon rank sum tests.

#### 2.5.2 fMRI

All preprocessing steps and subsequent analysis were performed using custom MATLAB scripts and SPM 12 (Statistical Parametric Mapping). Spatial preprocessing consisted of rigid co-registration to the animal’s own anatomical scan for better visualization of the results. Co-registered functional images were then resliced to 1 mm^3^ isotropic voxel size and smoothed with a Gaussian kernel (full width at half maximum: 1.5 mm). Single-subject analyses were performed using a fixed-effects general linear model to estimate the response amplitude at each voxel. The onset and duration of the stimulation blocks were used to generate a boxcar model, convolved with a MION hemodynamic response function [24]. The level of significance was set a p < 0.0001, uncorrected for multiple comparisons with a minimum cluster size of 20 [25,26]. The exact locations and extents of ECS-induced fMRI activations were verified on the animal’s own EPI images.

To evaluate the percent signal change (PSC) in the hippocampus, a functional ROI was created using half of the data (even runs), while the PSC was calculated on the other half (uneven runs) to avoid circularity. Standard error of the mean (SEM) was calculated over runs. Significant changes in PSC compared to the nonstimulation condition set as baseline were calculated using Wilcoxon signed-rank tests, Bonferroni-corrected for the number of ROIs. To examine the spread of ECS-induced activations in the hippocampus in a more detailed manner, we defined 9 seeds (1 mm radius spheres; spaced 2 mm apart) along the anterior-posterior axis of the hippocampus and 9 seeds in area TE along the same axis. PSC was calculated in the same manner as for anatomical and functional ROIs. The percentage activated voxels was calculated for both stimulation frequencies and tested using a Chi square test of independence or a Fisher exact test.

## 3 Results

Modelling of the electric field caused by 3 mA ECS showed the highest field strength in the cortex underneath the stimulation electrodes, at the level of the upper and lower banks of the Superior Temporal Sulcus, with a gradual decline in electric field strength from lateral to medial (Figure 1C). According to this model, the electric field would be minimal at the level of the hippocampus. In a separate experiment, we measured the electric field strength at 5 recording locations in depth and at two recording locations in the coronal plane (one at the level of lateral temporal cortex and one at the level of the hippocampus (Supplementary Table 1). The average electric field strength at the depth of the recording sites (last three measurements) was 5.3 V/m.

### 3.1 Behavioral effects of ECS on spatial memory

We investigated whether ECS could influence performance in a non-navigational spatial memory task. Each monkey underwent 7 sessions with stimulation at 40 Hz and 7 sessions at 10 Hz. ECS at 3 mA and 40 Hz (Figure 3A) significantly improved the overall performance on the stimulated patterns compared to the nonstimulated patterns in both monkeys (79.31% and 60.68% correct for stimulated patterns compared to 67.88% and 53.07% correct for nonstimulated patterns in monkey P and T, respectively; χ^2^ (1, N = 810) = 12.05, *p* = .0005 for monkey P and χ^2^ (1, N = 2329) = 12.87, *p* = .0003 in monkey T). Sham stimulation at 40 Hz (0.1 mA) had no effect (in monkey P, *p* = 0.06) and a very weak effect (monkey T, *p* = 0.02).

**Figure 3.**
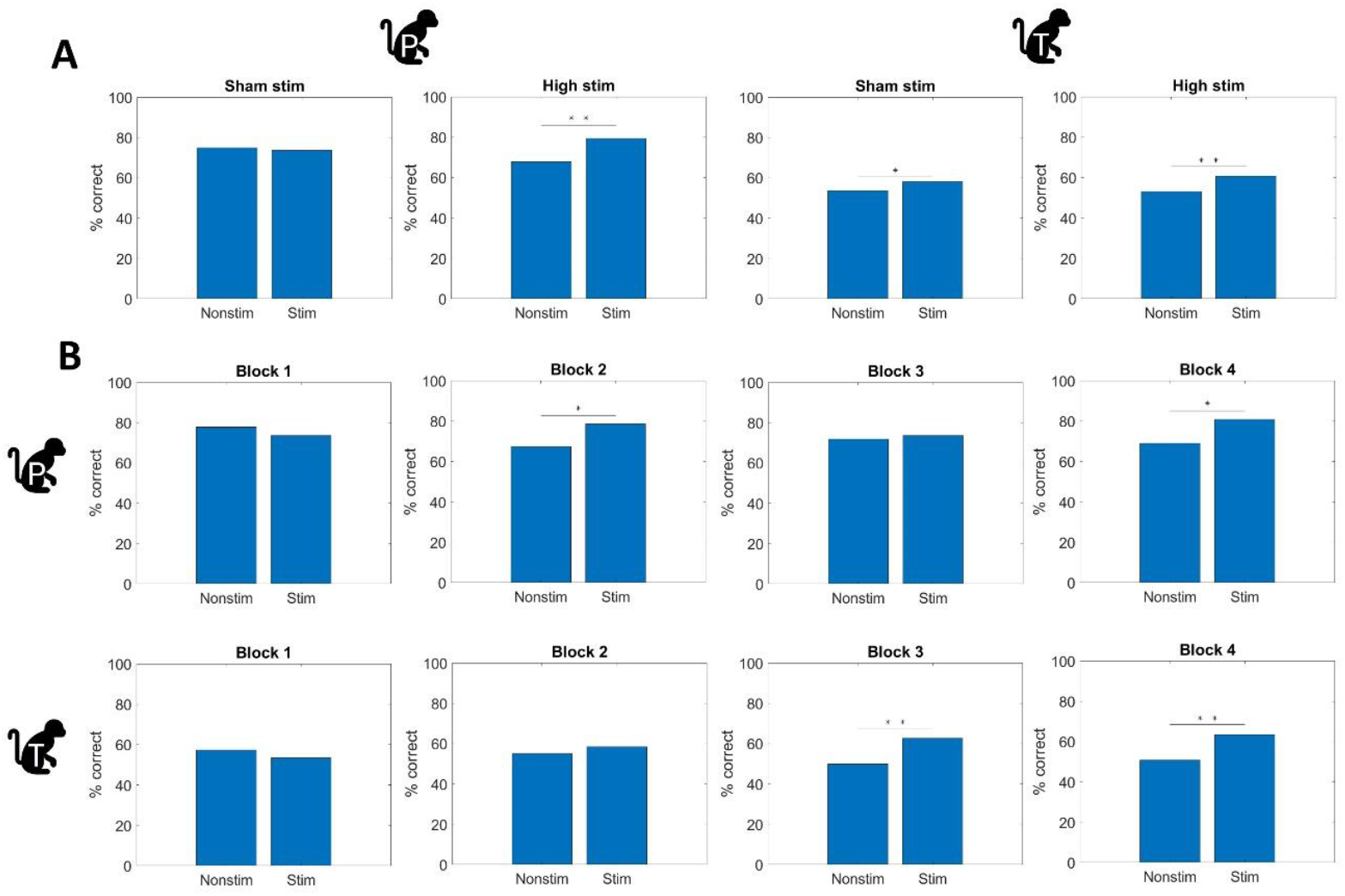
Behavioral effects of 40 Hz ECS. (A) Percent correct for nonstimulated and stimulated patterns during sham stimulation and high intensity (3 mA) stimulation for monkey P (left) and monkey T (right). (B) Percent correct for nonstimulated and stimulated patterns for each block separately for monkey P (top row) and monkey T (bottom row). Asterisks indicate statistical significance (Chi square test of independence,*: p < .05, **: p < .001).

We then determined whether ECS in a block of trials at 40 Hz would affect performance in a subsequent block of trials with sham stimulation (Figure 3B). During the first block (sham stimulation at 0.1 mA), there was no significant difference in performance between stimulated and nonstimulated patterns (Chi-square tests, Monkey T: χ^2^ (1, N = 1269) = 1.90, *p* = .168; Monkey P: χ^2^(1, N = 564) = 1.30, *p* = .255). In subsequent blocks, monkey P showed significant improvements in both high stimulation blocks (block 2: χ^2^(1, N = 581) = 8.13, *p* = .0043 and block 4: χ^2^(1, N = 229) = 3.91, *p* = .048), but not in the sham stimulation block 3 (χ^2^(1, N = 581) = 0.22, *p* = .641). However, Monkey T showed no significant difference between stimulated and nonstimulated patterns in the first high stimulation block (χ^2^(1, N = 1257) = 1.45, *p* = .229), but did perform better on the stimulated patterns in block 3 (sham stim) and 4 (high stim), resp. χ^2^(1, N = 1252) = 21.10, *p* < .001 and χ^2^(1, N = 1072) = 15.87, *p* < .001). To further validate the results of the Chi-square test, we fitted a GLM model. At 40 Hz, ECS caused a significant improvement across blocks, and this improvement became stronger with each block (Block 2: OR = 1.280, 95% CI [1.055, 1.553], Block 3: OR = 1.493, 95% CI [1.234, 1.806], and Block 4: OR = 1.718, 95% CI [1.360, 2.169]). At 10 Hz, ECS caused a significant reduction in Block 2 (high stim, OR = 0.778, 95% CI [0.633, 0.957]). This effect was not visible in Block 4 (high stim) due to the differing effects in the two monkeys. We also fitted the GLMM for each monkey separately, and the results matched those from the original Chi-square analysis. In the GLMM, a significant three-way interaction among the main effects indicated a block- and frequency-dependent effect of stimulation.

The behavioral results in monkey T were puzzling but could be explained by a partial spillover effect from block 2 (high stim) to block 3 (sham stim). To investigate the temporal dynamics of the ECS-induced behavioral improvement, we calculated running averages of the performance within each block. Monkey P consistently performed better on stimulated patterns during high stim blocks (Figure 4), and showed no difference in performance in block 3 (sham stim). In monkey T, the performance for both groups was similar in the first half of block 1 (the decrease in performance at the end of block 1 for the stimulated patterns was entirely due to a very low performance in session 1). In block 2 (high stim), performance remained similar for stimulated compared to nonstimulated patterns in the first half of the block, but improved for stimulated patterns in the second half of block 2 (Wilcoxon rank sum test on percentage correct of running average, W = 62.5, *p* = 0.042 on the last 9 data points). This improvement persisted but gradually declined in block 3 (Wilcoxon rank sum test between first half and second half of stimulation trials in block 3, W = 135, *p* = 0.04). Finally, the improved performance reappeared in block 4 and grew even stronger in the second half of that block, as evidenced by the significantly better performance for stimulated patterns in the second half of block 4 compared to the second half of block 2 (χ^2^(1, N = 452) = 4.25, *p* = .039). Nonstimulated pattern performance fluctuated within blocks but remained relatively constant overall. Monkey T consistently responded faster after stimulated patterns in high-intensity ECS blocks (Wilcoxon rank sum test; z = 4.74, p = 0.00002 and z = 5.35, p < 0.00001 in block 2 and 4, respectively) but not in sham stimulation blocks (Figure S1). Thus, the lack of a significant effect on performance in block 2 in this animal may also be explained by a speed-accuracy tradeoff in this high-intensity stimulation block. Monkey P did not show any consistent difference in reaction time between stimulated and nonstimulated patterns (with the exception of a marginally significant difference in block 3). We also plotted the mean percentage correct in the first sham block of each sessions (Figure S2). Day-to-day performance was variable, and no systemic difference between the nonstimulated and stimulated patterns was observed, thus there does not seem to be a clear effect of memorization or stimulation across days.

**Figure 4.**
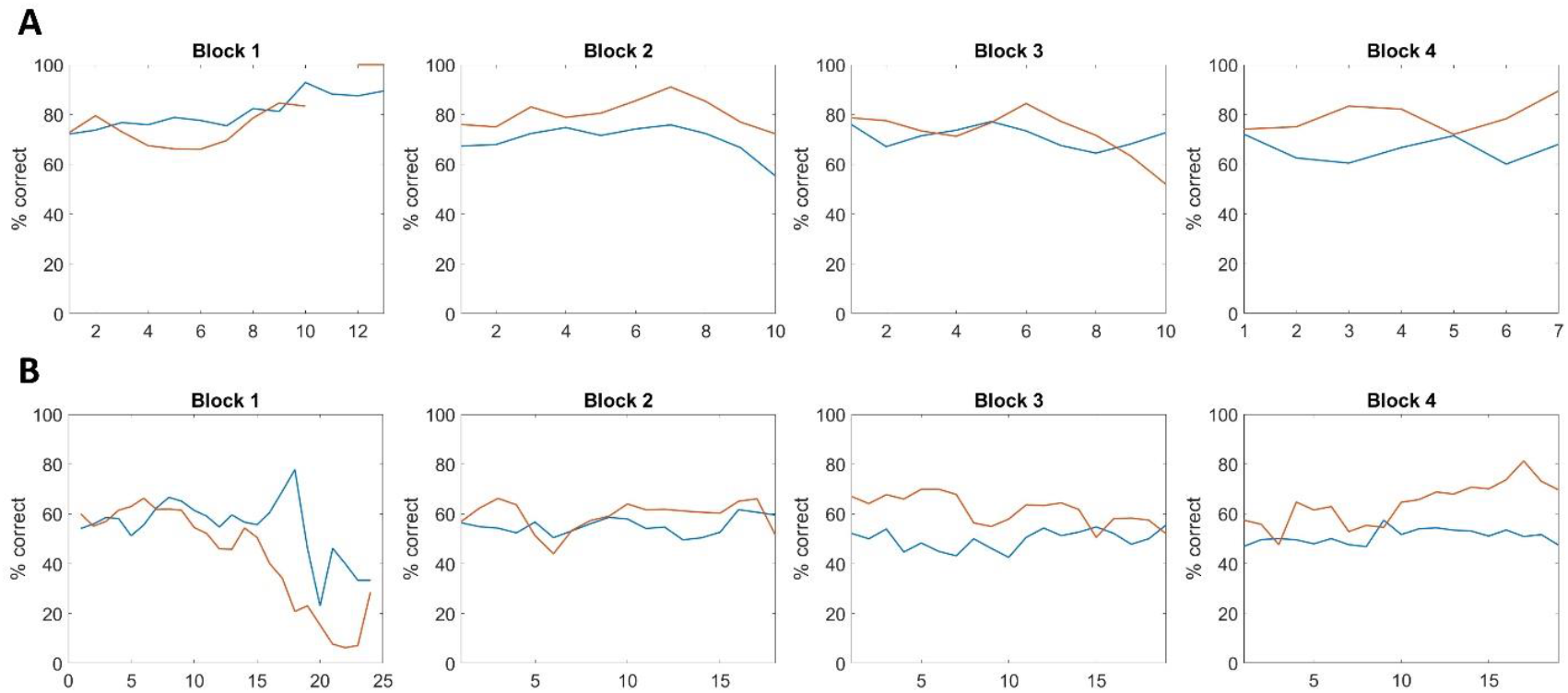
Evolution of performance during 40 Hz ECS sham (blocks 1 and 3) and high intensity (blocks 2 and 4). (A) Running average of the mean performance for nonstimulated (blue) and stimulated patterns (red) for monkey P. For each data point, the mean of 25 trials was calculated with a sliding window of 10 trials. ECS was applied at 3 mA 40 Hz in the high stim blocks. (B) Same data for monkey T.

As a control, we also recorded 7 sessions with ECS at 10 Hz following the same paradigm as the 40 Hz ECS (Figure S3 and S4). Monkey P showed small and inconsistent effects of high-intensity 10 Hz ECS (a marginally significant decrease in performance for stimulated patterns in block 2, χ^2^(1, N = 707) = 4.20, *p* = .040 and a weak increase in performance in block 4, χ^2^(1, N = 623) = 6.05, *p* = .014). However, monkey T showed a highly significant decrease in performance for stimulated patterns at high-intensity 10 Hz ECS in block 3 (χ^2^(1, N = 1087) = 10.71, *p* = .001) and block 4 (χ^2^(1, N = 997) = 11.97, *p* < .0001). For reaction times during 10 Hz ECS, see Figure S5.

Additionally, we recorded four sessions in monkey P in which we stimulated at 80 Hz (see Figure S6). We fitted a GLMM that included the first four sessions of 40 Hz data and the four sessions at 80 Hz. There was no significant effect of stimulation in the 80 Hz ECS blocks (Block 1: OR = 0.824, 95% CI [0.526, 1.290], Block 2: OR = 0.753, 95% CI [0.425, 1.335], Block 3: OR = 1.117, 95% CI [0.661, 1.888], and Block 4: OR = 0.816, 95% CI [0.499, 1.334]). At 40 Hz, we observed the same results as when including all seven sessions.

### 3.2 Widespread functional activations caused by ECS

To examine the effects of ECS across the brain, we performed ECS at 40 Hz and 10 Hz in two sedated monkeys in a block-design fMRI experiment. ECS at 40 Hz induced an extensive pattern of activations and deactivations throughout the brain, whereas 10 Hz ECS elicited very few activations and deactivations (Figure 5 and Supplementary Figure S7). Both the number of significantly activated voxels and the number of significantly deactivated voxels was markedly higher for 40 Hz ECS than for 10 Hz ECS in both monkeys (Monkey C: activations, χ^2^(1, N = 97898) = 4378.92, *p* < .0001; deactivations, χ^2^(1, N = 97898) = 5242.38, *p* < .0001; Monkey P: activations, χ^2^(1, N = 96023) = 6701.61, *p* < .0001; deactivations χ^2^(1, N = 96023) = 8690.36, *p* < .0001).

**Figure 5.**
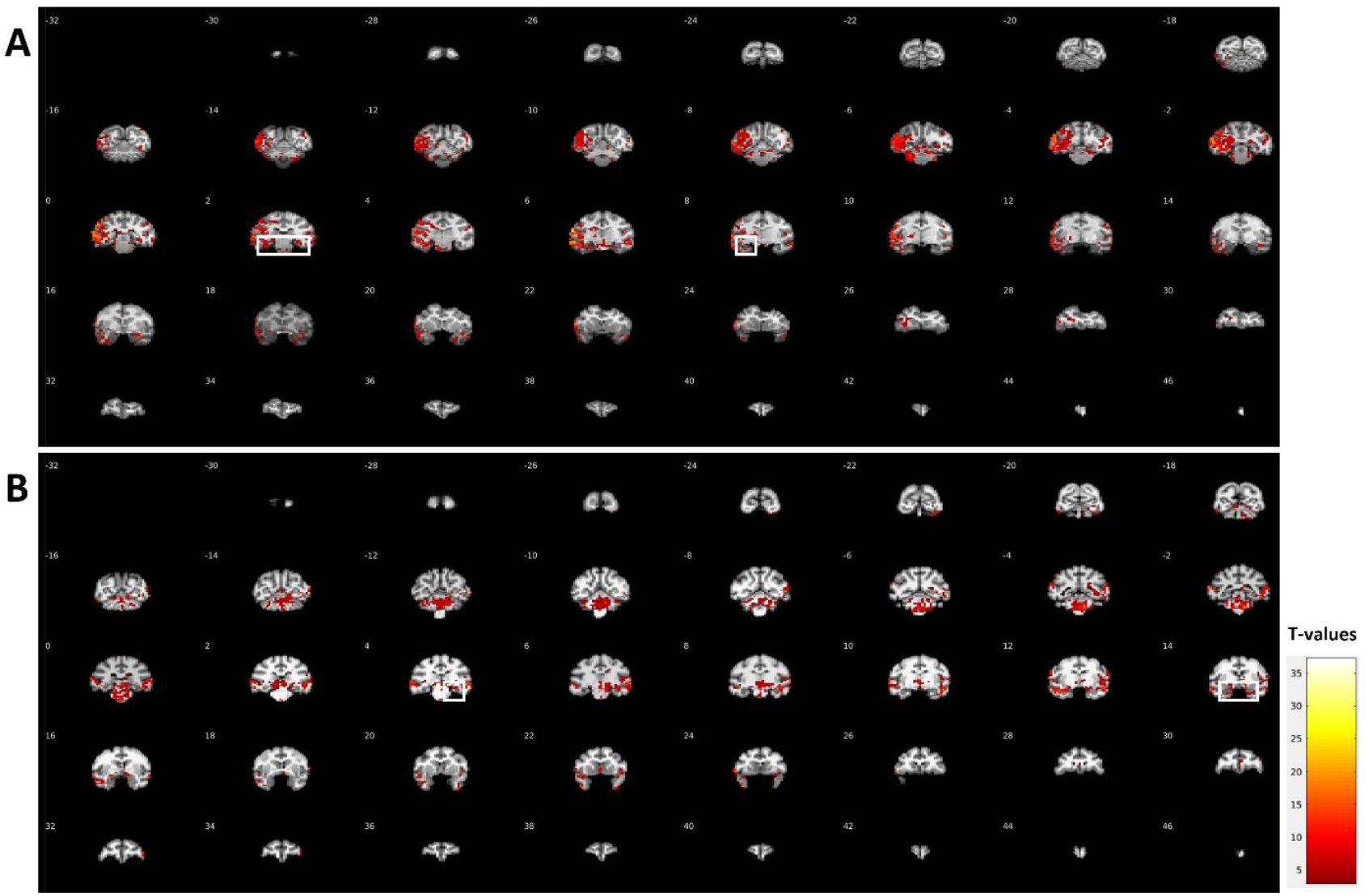
Overview of fMRI activations during 40 Hz ECS. (A) Data for Monkey P. T-score maps of the contrast stimulation versus baseline (uncorrected p < 0.0001) overlaid on the monkey’s own anatomical MRI. White squares highlight activations in anterior and posterior hippocampus. (B) Data for Monkey C. Same conventions as in (A).

In both monkeys, ECS at 40 Hz strongly activated parts of the hippocampus and deactivated other parts, but 10 Hz ECS did not (example sections in Figure 6A and B). We quantified the PSC within anatomically defined hippocampal ROIs for each monkey (Figure S8). At 40 Hz ECS, the PSC was increased significantly in the left hippocampus of monkey P, while the PSC was decreased non-significantly in the left hippocampus of monkey P and in both hippocampi of monkey C. At 10 Hz, the PSC was decreased in three out of four hippocampi, but none of these effects reached significance. Overall, the anatomical ROI analysis was dominated by inactivation and likely underestimated spatially focal responses; therefore, subsequent analyses used a functionally defined ROI in the hippocampus based on half of the runs of the 40 Hz data, and we calculated the PSC using the other half of the runs. The PSC in this functionally defined ROI was significantly higher than zero in all 4 hemispheres, whereas 10 Hz ECS did not induce any hippocampal activation (Figure 6C). Moreover, the number of activated voxels in an anatomical hippocampal ROI based on the monkeys’ own MRI differed significantly between the two frequencies in both monkeys (Fisher’s exact test, *p* < .00001, for both the bilateral hippocampus and for the two hemispheres separately).

**Figure 6.**
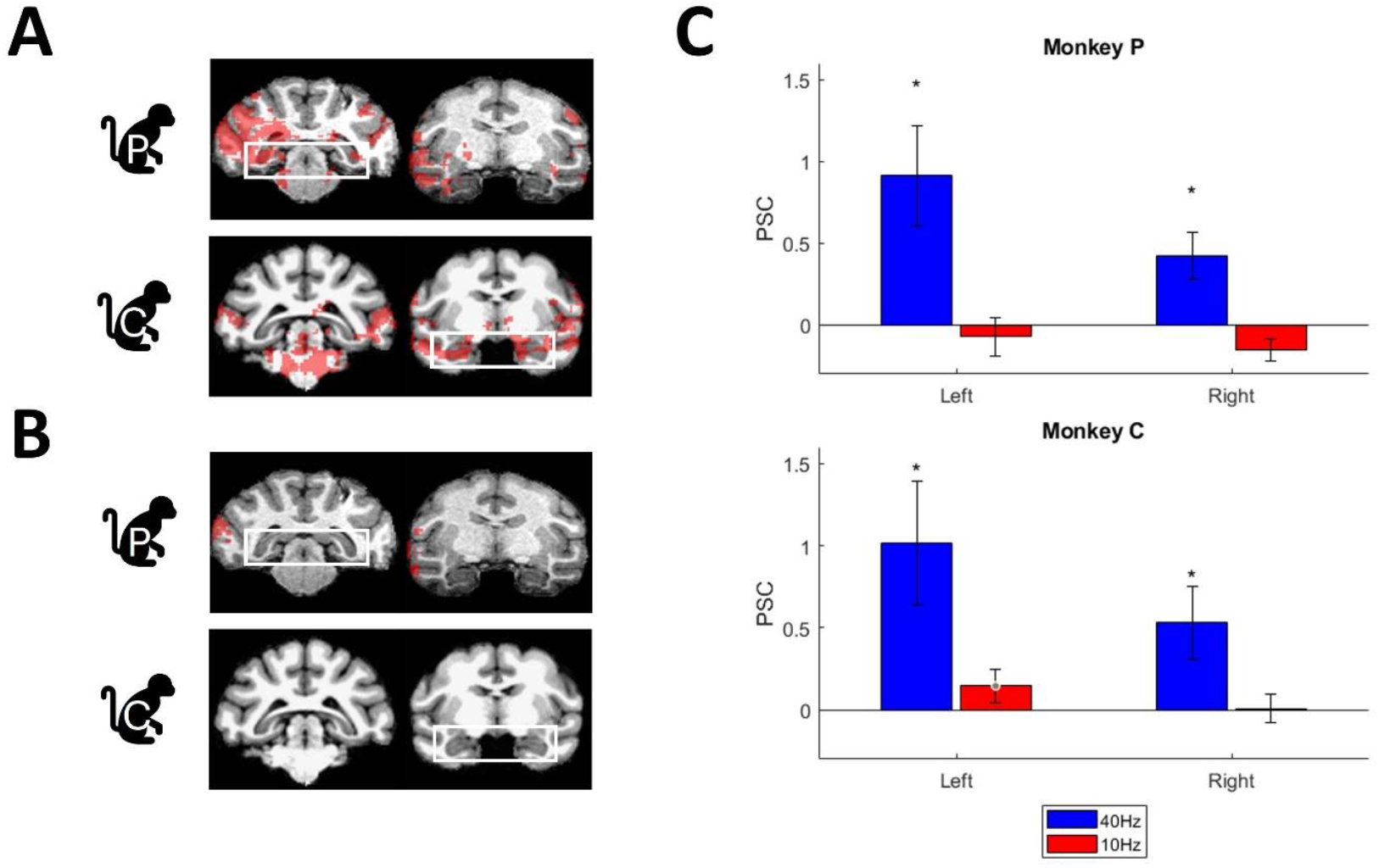
Comparison of the fMRI activations at 40 Hz and 10 Hz. (A) Example sections from T-score maps of the contrast stimulation versus baseline at 40 Hz ECS (uncorrected p < 0.0001, cluster correction: 20). White boxes show the activations on which the functional ROI was defined to calculate the PSC. (B) The same sections as in A) at 10 Hz ECS. (C) Bar plots of percent signal change (PSC) in functional ROIs in the hippocampus for 40 Hz ECS-fMRI (blue) and 10 Hz ECS-fMRI (red). Top row shows results from monkey P and bottom row shows results from monkey C.

Figure 6A and B illustrate that monkey C mainly showed activations in the anterior hippocampus, whereas monkey P mainly showed posterior hippocampus activations. To visualize this effect, we plotted the PSC in seeds defined along the anterior-posterior extent of the hippocampus and in the more laterally located area TE, part of the inferotemporal cortex (Figure 7A). In monkey P, the PSC was significantly positive compared to nonstimulation blocks in the posterior hippocampus in both hemispheres but not more anteriorly. Conversely in monkey C, the PSC was predominantly negative in the posterior to mid-hippocampus, while the most anterior part of the hippocampus showed a significantly positive PSC (Figure 7A). ECS induced mostly bilateral hippocampal activations. Monkey C showed strong bilateral activations in the anterior hippocampus (Figure 6A), and an additional activation in the right hemisphere at the level of the mid-hippocampus. Similarly, monkey P also had an activation cluster in the anterior left hippocampus (data not shown). Despite these activation clusters, the overall PSC in the anatomically defined ROIs in the hippocampus was mostly negative due to strong deactivations surrounding these clusters (Figure 7B). For the seeds defined in area TE, PSCs were significantly elevated posteriorly and decreased more anteriorly, except for the right hemisphere of monkey P (Figure 7A). The electric field modeling predicted the highest field strength in the lateral temporal cortex and minimal in the rest of the brain (Figure 1C). However, our fMRI data revealed strong activations not only in lateral temporal cortex but also in multiple other regions (Figure S9), indicating that the electric field may be more widespread than predicted. Notably, we observed the strongest hippocampal activations at anterior-posterior levels where no significant TE activations were present, which suggests that at least part of the hippocampal activations were frequency-specific. Effect size for all PSCs calculations can be found in the Supplementary Data (Table S3 and S4). Seed-based functional connectivity analysis using either anatomical or functional hippocampal ROIs failed to show any significant changes with TE (data not shown).

**Figure 7.**
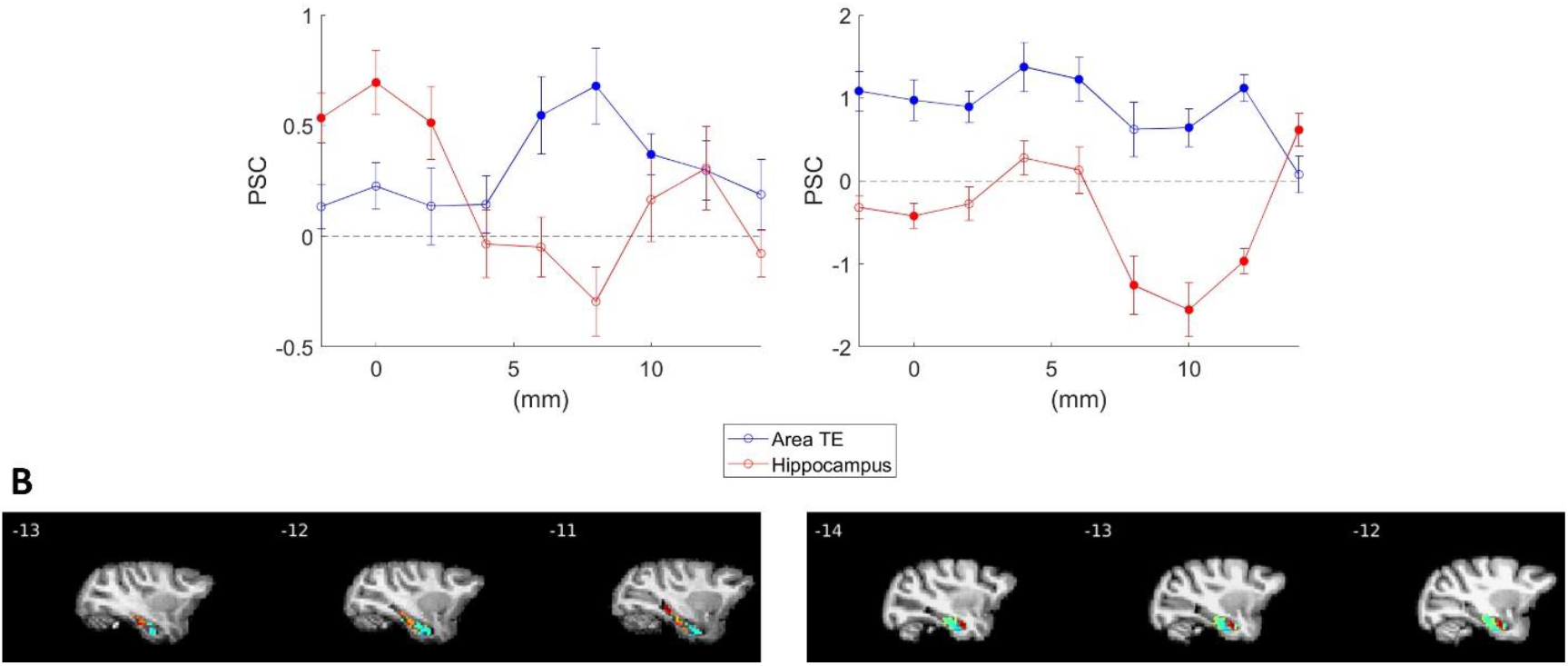
Seed-based analysis of 40 Hz ECS-fMRI activations. (A) Percent signal change in seeds defined along the posterior-anterior axis in the hippocampus (red) and in area TE (blue) of the right hemisphere in monkey P (left) and monkey C (right). Filled symbols indicate significantly activated or inactivated seeds. Error bars indicate standard error of the mean. Significance was tested by a two-sided Wilcoxon signed rank test: * p < 0.05; corrected for multiple comparisons. (B) T-score maps (sagittal sections) of the contrast stimulation versus baseline (uncorrected p < 0.0001, cluster correction: 20) overlaid on the monkey’s own anatomical template (monkey P: left, monkey C: right). A mask was used to only include the hippocampus to increase visibility.

For activations in other cortical anatomical ROIs see Figure S9 and for subcortical activations see Figure S10.

## 4 Discussion

Epicranial current stimulation (ECS) at 40 Hz significantly improved performance in a spatial memory task in both animals, whereas ECS at 10 Hz did not. ECS at 40 Hz – but not at 10 Hz – also induced widespread fMRI activations in cortical and subcortical regions including the hippocampus. Overall, our findings reveal the potential of ECS at 40 Hz as a minimally-invasive approach to improve spatial memory.

Forty Hz stimulation is capable of engaging the hippocampus, when applied through both electrical or sensory stimulation [20,27–29]. Gamma oscillations, along with theta oscillations, are the primary rhythms present in the hippocampus. These gamma oscillations originate from two independent systems: one located in the entorhinal cortex and one in CA3. The primary mechanism involves rhythmic inhibitory postsynaptic potentials (IPSPs) in the pyramidal cells, supported by fast-spiking interneurons that drive inhibition in these cells [16–18]. Consequently, applying 40 Hz electrical stimulation can enhance the periodic firing of these interneurons, leading to increased gamma entrainment and facilitating memory encoding and retrieval [17]. Besides an effect on the hippocampus itself, ECS can also increase the functional connectivity between the hippocampus and key brain regions and networks such as the posterior inferior parietal cortex and posterior medial network [30–32]. The posterior parietal cortex (PPC) and dorsolateral prefrontal cortex (dlPFC) are the main brain regions involved in spatial working memory [33]. Our fMRI results did not show any dlPFC activation in both monkeys and only minimal activation in the PPC in one monkey. Nonetheless, the functional connectivity during a spatial memory task between the hippocampus and PPC and/or dlPFC could still be enhanced as our fMRI was performed in sedated monkeys.

ECS at 40 Hz enhanced memory performance in both animals, but the timing of this improvement varied between the two monkeys. One animal exhibited enhanced memory performance exclusively during ECS, while the other animal showed a delayed effect of stimulation that persisted during subsequent blocks. This delayed effect may be attributed to a partial spillover from the prior high stimulation block. A meta-analysis on the effectiveness of tACS for improving cognition in humans found that offline improvements in performance were generally stronger than online improvements [10]. Therefore, it may not be surprising that we observed significant effects both during and after the application of ECS in this study. It is important to note that the observed performance improvements were not sustained beyond the stimulation session, as performance between nonstimulated and stimulated patterns was not significantly different in the first block the following day. Furthermore, tACS effects may arise from entrainment of the ongoing neural oscillations to the applied stimulation frequency or from modulation of neuronal firing timing, which could lead to spike-timing dependent plasticity (STDP). Given that STDP results from cumulative changes in neural activity over time, this could account for the spillover effect observed in our behavioral study [10,34,35]. Although measuring neural activity during electrical stimulation is challenging, future studies should investigate the effects of 40 Hz ECS at the level of individual neurons.

We employed a non-navigational spatial memory task, based on Forcelli et al. (2014), who demonstrated a causal role of the hippocampus in this type of task using muscimol inactivation [22]. Although the role of the hippocampus in navigation is well-established, some debate remains about its involvement in non-navigational spatial memory. Hippocampal lesions studies in monkeys often show no or minimal impairment in non-navigational spatial tasks [36,37]. These tasks can be solved using either allocentric (viewpoint-independent) or egocentric (viewpoint-dependent) cues [22]. While allocentric spatial processing is highly hippocampal dependent, evidence for hippocampal involvement in purely egocentric processing is limited [38]. The preserved spatial processing ability in hippocampal-lesioned monkeys over short delays may be explained by their reliance on intact egocentric spatial representations, supported by posterior parietal cortex [39,40]. However, at longer delays, these egocentric representations are no longer maintained, leading to decreased performance [39]. Nonetheless, some studies have shown a role for the hippocampus in certain forms of egocentric spatial processing [41–43]. In our study, the monkey was seated in the same position for every session in a dark room, which promoted the use of an egocentric strategy to solve the task. However, as the monkey was required to remember the location of two targets, there is a sequential component in the task, rendering the task more hippocampal dependent. A previous study found that patients with hippocampal lesions were not impaired in single memory-guided saccades, but were impaired in sequences of memory-guided saccades [44]. Future studies should evaluate the efficacy of 40 Hz ECS on different memory tasks to determine whether its effects generalize to other forms of memory.

ECS at 10 Hz did not consistently improve behavioral performance in our monkeys. Increased alpha activity over visual processing regions can be associated with impaired visual perception, thereby reducing the processing of the target locations [45]. Reduction in alpha power over the contralateral hemisphere occurs in covert spatial attention studies in humans (EEG) [46]. However as we stimulated bilaterally in our experiment, alpha power was most likely increased over both hemispheres, potentially counteracting the reduction of posterior alpha power and leading to inconsistent memory effects.

During ECS-fMRI, we found activations in both hippocampi in two animals and widespread activations across the brain. Although the number of concurrent tACS-fMRI studies is limited, previous research indicates that the strongest activations often occur outside the regions directly beneath the stimulation electrodes. Similarly, intracranial recordings in rats found neural activations distant from the stimulation electrodes, suggesting that tES activates neurons both directly and indirectly via polysynaptic connections [47,48].

We observed a highly significant difference in the number of activations caused by 40 Hz and 10 Hz ECS during fMRI. This effect could be due to the fact that 40 Hz stimulation delivers four times the amount of current compared to 10 Hz when applied with the same duration. However, previous studies have suggested that tES induces frequency-specific effects due to the differential susceptibility of brain regions to entrain to specific frequencies [48,49]. Our seed-based analysis in the hippocampus and area TE revealed largely dissociated activation patterns along the anterior-posterior axis. Despite the strongest electric field occurring in the cortex beneath the stimulation electrodes, we observed significant activations in the hippocampus at anterior-posterior levels where no activations in area TE were observed, and vice versa. This suggests that our hippocampal activations were not merely resulting from a simple electric field gradient induced by ECS [58,59]. Our behavioral results further support our hypothesis that our results were frequency specific as 80 Hz ECS did not cause any effect in performance while 40 Hz ECS did. If these results were purely dose-dependent, we would have expected a behavioral effect as well.

Our ECS-fMRI study was performed under sedation because performing the spatial memory task in the scanner would have induced large artefacts caused by the arm movements arm. A previous study using intracortical microstimulation in parietal cortex during fMRI has demonstrated highly comparable functional activations in awake animals compared to during ketamine sedation [25]. Although the behavioral results and the fMRI data were collected in different states (awake vs ketamine sedation), the selective hippocampal activations at 40 Hz may have been responsible for the behavioral improvement we observed. Finally, although the exact location of the epicranial electrodes differed slightly between the subjects, memory improvement during 40 Hz ECS was observed in both subjects as well as the hippocampal activations during fMRI. This finding suggests that the frequency of ECS is more important than the exact implantation location of the electrodes.

Entorhinal DBS in humans improves spatial and recognition memory, but direct hippocampal stimulation did not affect performance [7,8]. The possibility exists that our ECS stimulated entorhinal white matter tracts projecting to the hippocampus, thereby causing the hippocampal activations and memory improvement. ECS may have the additional advantage over DBS that the induced electric field is most likely much wider compared to the one induced by DBS on a single electrode. A minimally-invasive, safe and effective approach for stimulation-induced memory improvement may represent a major breakthrough for patients suffering from memory impairment.

## 5 Conflict of interest declaration

The authors declare that they have no known competing financial interests or personal relationships that could have appeared to influence the work reported in this paper.

## 6 Acknowledgements

Funded by Mission Lucidity, Fonds voor wetenschappelijk onderzoek Vlaanderen (G. G0D6520N) and KU Leuven grant C14/22/134 and TOP/22/019. We thank Nick Van Helleputte for helpful discussions and Inez Puttemans, Christophe Ulens, Astrid Hermans, Stijn Verstraeten, Wouter Depuydt and Marc Depaep for technical assistance.

## Notes

### Competing Interest Statement

The authors have declared no competing interest.

