## Supplementary figures for "Epicranial electrical stimulation improves non-navigational spatial memory in macaque monkeys"

| Depth | 1 | 2 | 3 | 4 | 5 |
|---|---|---|---|---|---|
| Electric field (V/m) | 0.2 | 2.85 | 5.05 | 4.5 | 6.35 |

**Table S1. Electric field values** in V/m at 5 recording depths. Depth 1 corresponds to the top part of the hippocampus.

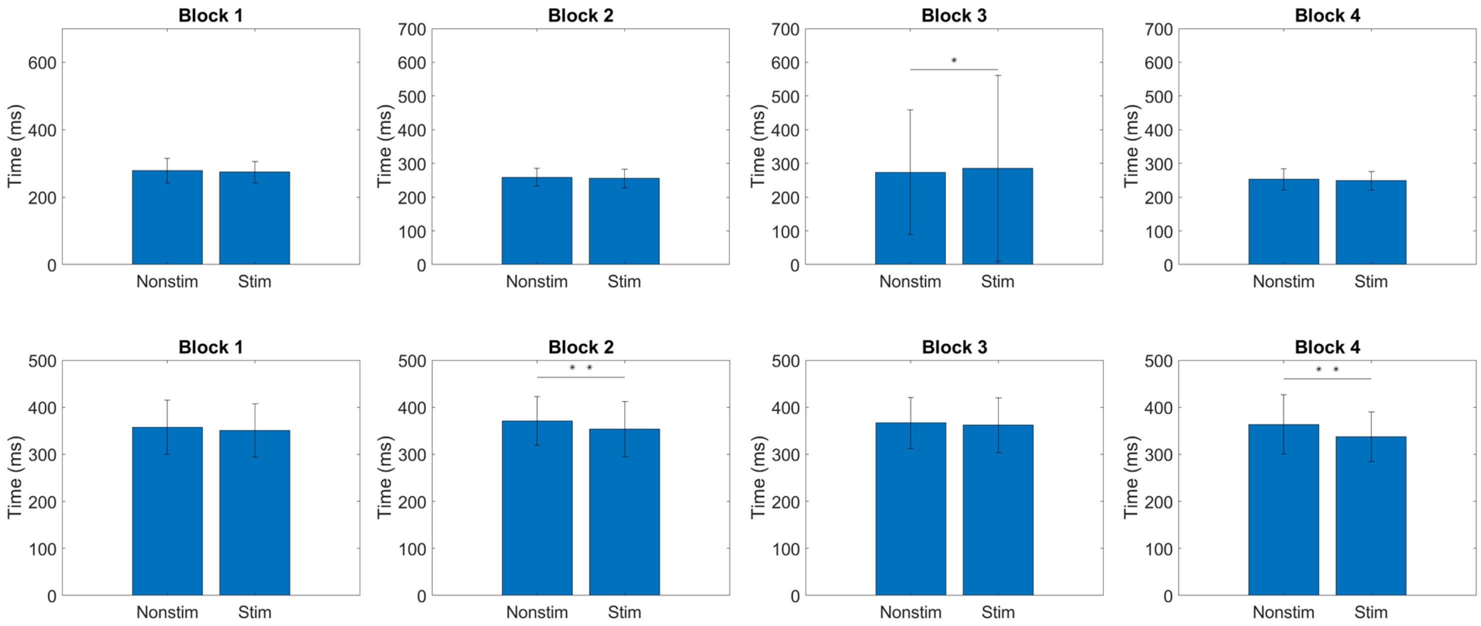


**Figure S1. Reaction times 40 Hz ECS.** Bar plots showing mean reaction time for nonstimulated versus stimulated patterns for monkey P (top row) and monkey T (bottom row). Asterisks indicate statistical significance (Wilcoxon rank sum test: *, $p < 0.05$). ECS was applied at 3 mA 40 Hz in the high stim blocks.

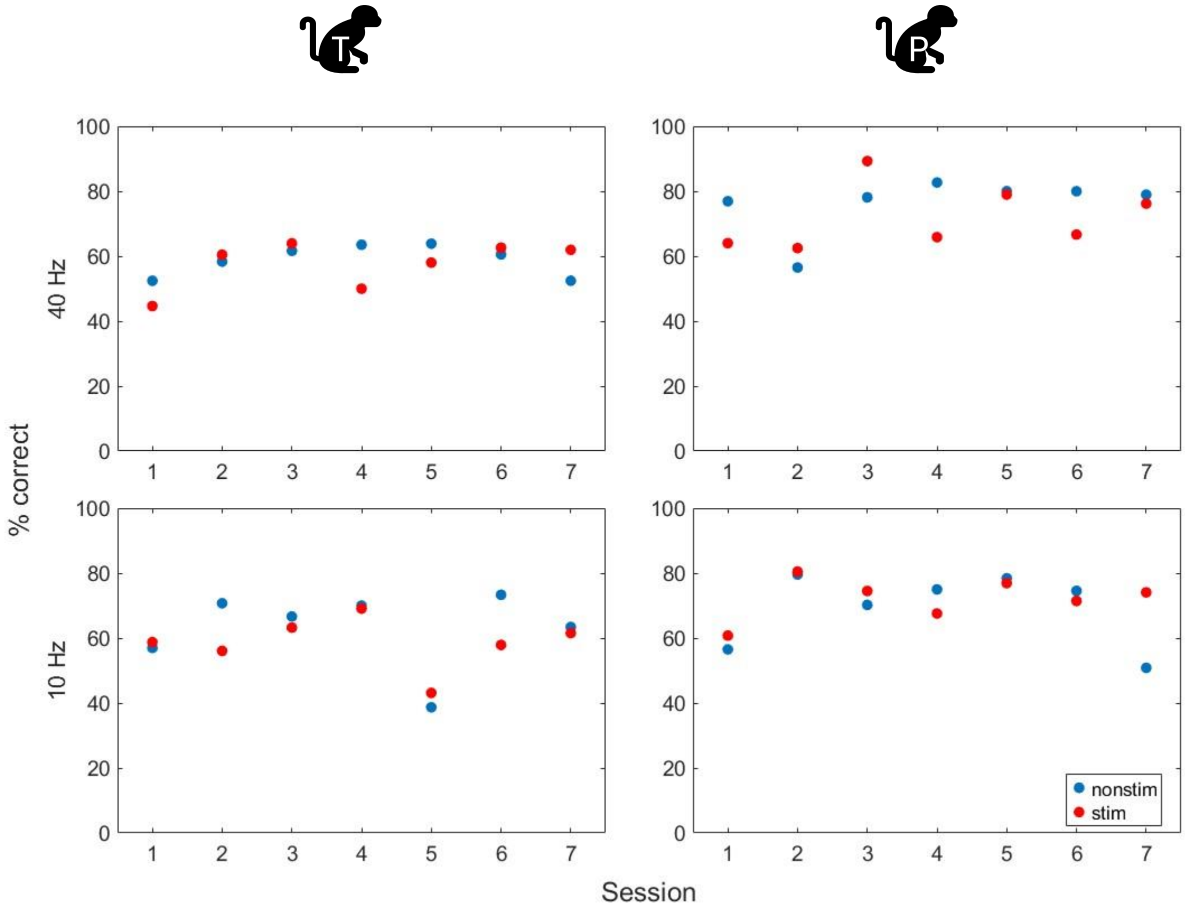


**Figure S2 Mean performance in the first sham block of each session** for 40 Hz ECS (top row) and 10 Hz ECS (bottom row). Blue dots show the mean percentage correct for nonstimulated patterns and red dots for stimulated patterns for monkey T (left) and monkey P (right).

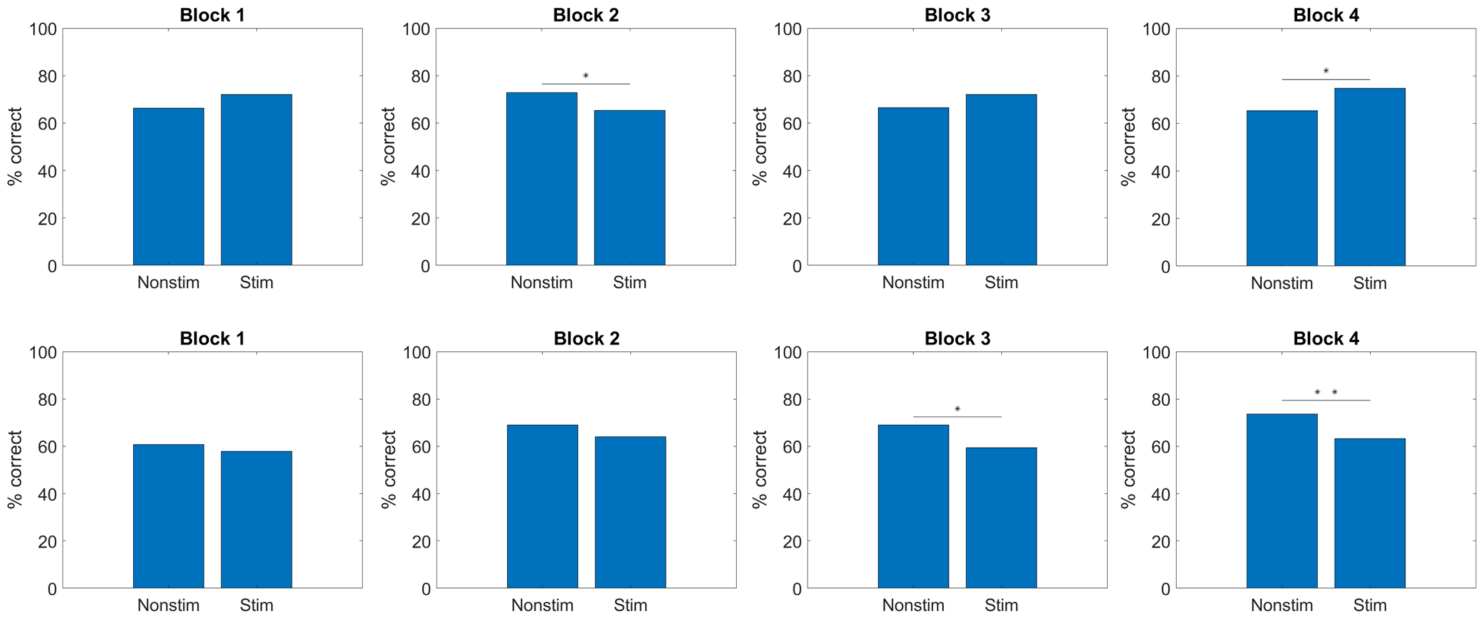


**Figure S3. Behavioral effects of 10 Hz ECS.** Percent correct for nonstimulated and stimulated patterns for sham (blocks 1 and 3) and high intensity (3 mA, blocks 2 and 4) stimulation for monkey P (top row) and monkey T (bottom row). Asterisks indicate statistical significance (Chi square test of independence, *: $p < .05$, **: $p < .001$).

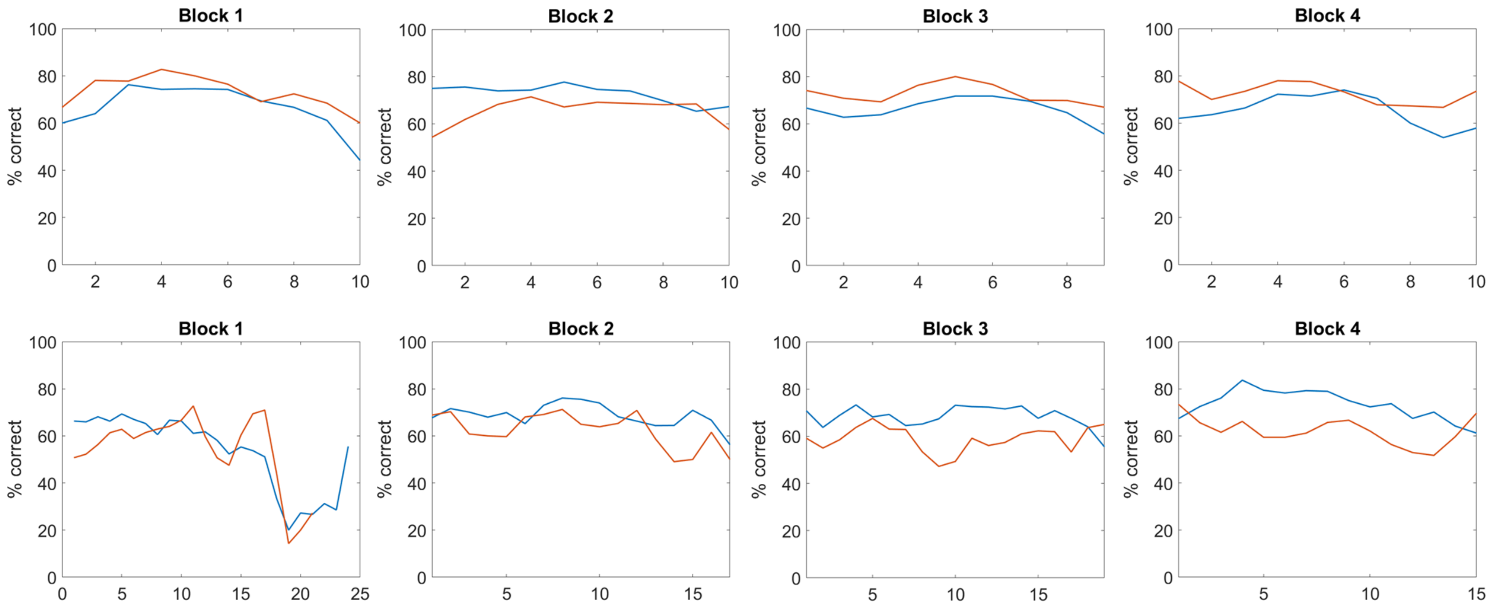


**Figure S4. Evolution of performance during 10 Hz ECS sham (blocks 1 and 3) and high intensity (blocks 2 and 4).** Running average of the mean performance for nonstimulated (blue) and stimulated patterns (red) for monkey P (top row) and monkey T (bottom row). For each data point, the mean of 25 trials was calculated with a sliding window of 10 trials. ECS was applied at 3 mA 10 Hz in the high stim blocks. Analogous to the 40 Hz experiment, the performance on stimulated patterns decreased already in the second half of block 2 (Wilcoxon rank sum test on percentage correct of running average, W = 88, $p$ = 0.0362 on the last 8 data points) and persisted in block 3. In additional sessions (not included in this behavioral study), the monkey even stopped responding to stimulated patterns after 10 Hz ECS at 3 mA. When stimulation intensity was set at 0.1 mA, however, the monkey responded again to these patterns.

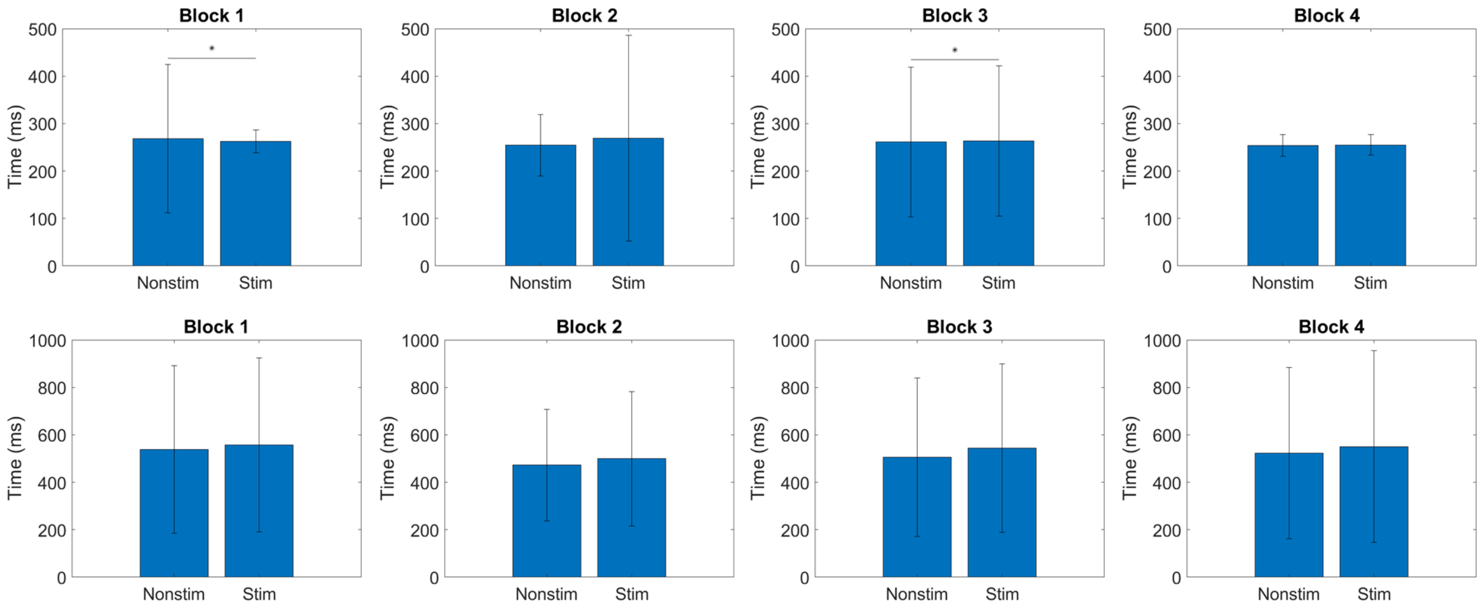


**Figure S5. Reaction times 10 Hz ECS.** In the 10 Hz sessions, reaction times did not differ significantly between nonstimulated and stimulated patterns for monkey T (bottom row) across all blocks (Supplementary figure 2). For monkey P (top row), reaction times for stimulated patterns were significantly higher compared to nonstimulated patterns but this difference was already present in the first block (Wilcoxon rank sum test, Block 1: $z = -2.79$, $p = 0.005$ and Block 3: $z = -2.38$, $p = 0.017$). The total time required to execute a trial (i.e. the time from the go-cue until touching the second target position) was significantly higher for stimulated patterns than for nonstimulated patterns in monkey T (suggesting that the animal took more time to compensate for the effect on accuracy), but lower for stimulated patterns in monkey P.

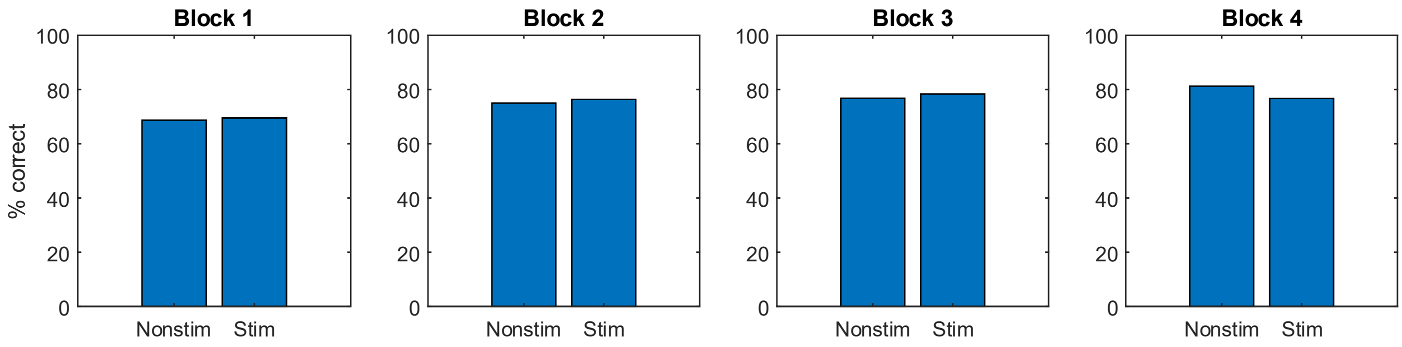


**Figure S6 Behavioral results at 80 Hz ECS.** Percent correct for nonstimulated and stimulated patterns for sham (blocks 1 and 3) and high intensity (3 mA, blocks 2 and 4) stimulation for monkey P.

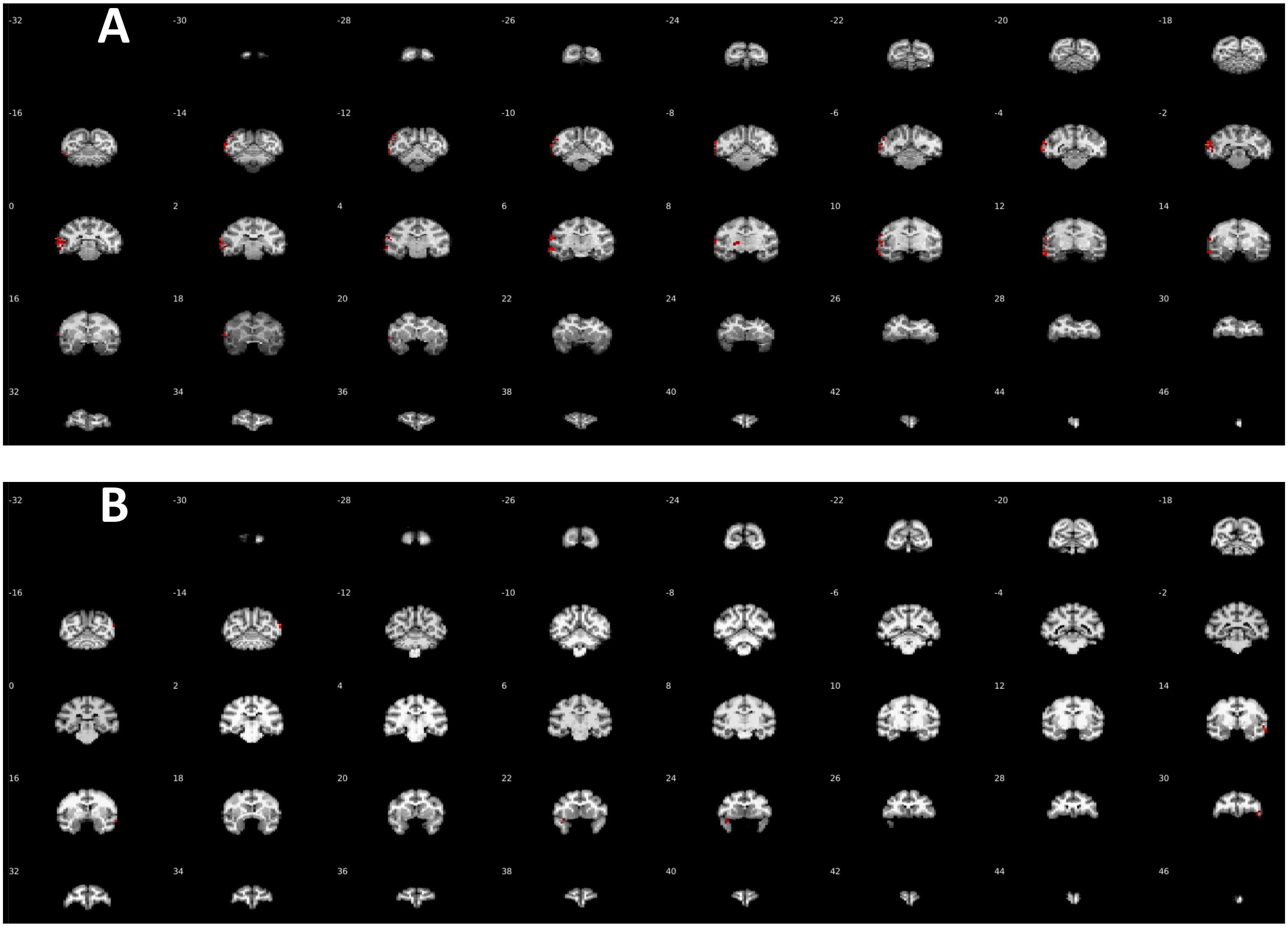


**Figure S7. Overview of activations in 10 Hz ECS-fMRI** (A) Monkey P. T-score maps of the stimulation frequency versus baseline (uncorrected $p < 0.0001$, cluster correction: 20) overlaid on the monkey's own anatomical template. (B) Monkey C. Same conventions as in (A)

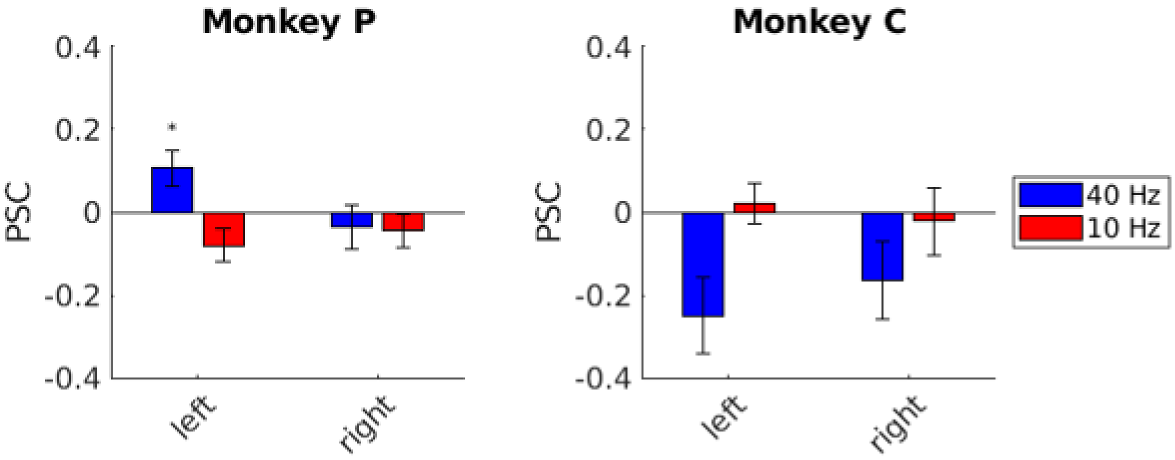


**Figure S8 PSC of the anatomical ROIs of the hippocampus** for monkey P (left) and monkey C (right). Asterisks indicate significance at $p < 0.05$, corrected for multiple comparisons.

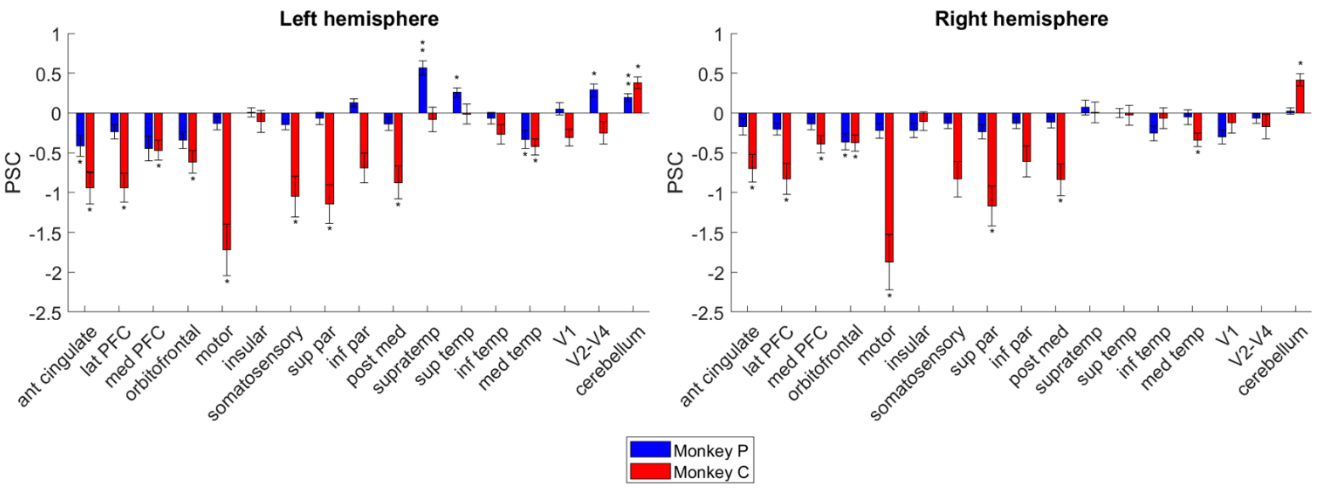


| **Table 2. Overview of anatomical ROIs based on CHARM atlas and Paxinos** | | |
|---|---|---|
| *Frontal cortex* | | |
| | Anterior cingulate | 24 32 |
| | Lateral prefrontal | 10 9 46 47 45 8 |
| | Medial prefrontal | 32 25 |
| | Orbitofrontal | 11 13 14 |
| | Motor | 4 6 44 |
| *Insular and gustatory cortex* | | |
| *Parietal cortex* | | |
| | Somatosensory | |
| | Superior parietal | PE Pea PECg PGM POM |
| | Inferior parietal | POa PO OPT PG |
| | Posteromedial | 23 29/30 7m 31 |
| *Temporal cortex* | | |
| | Supratemporal | PaAR AKM/AKL PaAC ST1 ST2 ST3 PaAL TPt Pa1 PrOK Rel |
| | Superior temporal | TAa TPO Pga/MST IPA FST TEa MT MTC TEM |
| | Inferotemporal | TE1 TE2 TE3 TEO |
| | Medial temporal | Presubiculum entorhinal perirhinal |
| *Occipital cortex* | | |
| Cerebellum | | |

**Figure S9. Overview of PSC during 40 Hz ECS in anatomical ROIs.** Average PSC (+/- SEM) per subject over runs for each hemisphere separately. Significance was tested by a two-sided Wilcoxon signed rank test: * $p < 0.05$; ** $p < 0.001$; corrected for multiple comparisons (17 ROIs). To quantify the 40 Hz ECS-induced effects in the rest of the brain, we defined 17 cortical anatomical ROIs and calculated PSC in each anatomical ROI. Although we measured significant functional activations in several cortical areas in both animals (e.g. in motor cortex, inferior parietal lobule, somatosensory cortex, supratemporal plane, superior temporal cortex, inferotemporal cortex, medial temporal lobe, occipital cortex, and the cerebellum, see also Figure 7), the simultaneous presence of strong deactivations resulted in predominantly negative PSCs in almost all anatomically-defined ROIs. Percent signal change (PSC) was calculated using MarsBaR (version 0.44) and averaged over runs. Anatomical regions of interest (ROI) were based on the Cortical Hierarchy Atlas of the Rhesus Macaque (CHARM, level 2) and Paxinos (see Table 2 for detailed description of ROIs; numbers correspond to Brodmann areas).

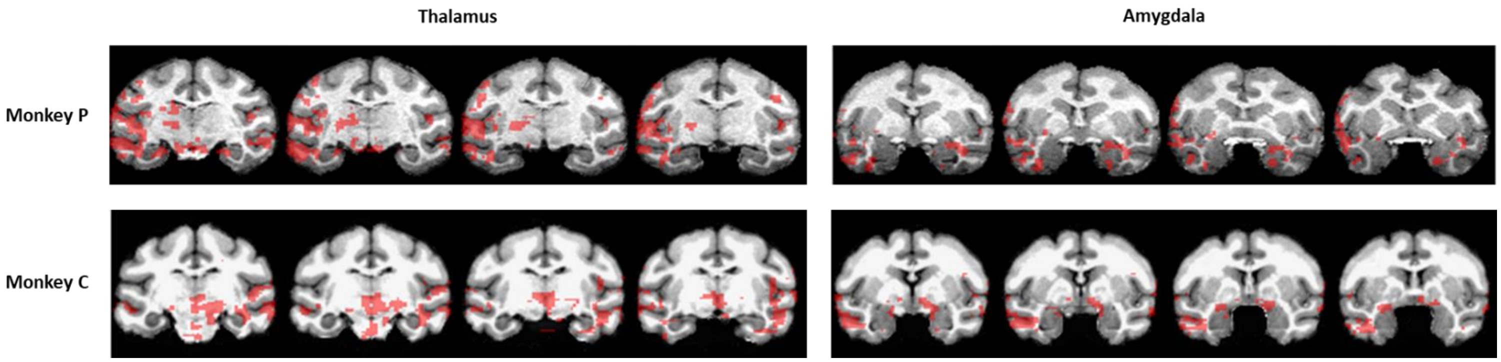


**Figure S10. Close-up of subcortical activations of 40 Hz ECS.** T-score maps of the stimulation frequency versus baseline (uncorrected $p < 0.0001$, cluster correction: 20) overlaid on the monkey's own anatomical template.. Monkey C had bilateral activations in the amygdala and in the thalamus, while monkey P had large activations in the right amygdala and in the left thalamus. Additionally, smaller activation clusters were present in the putamen, internal globus pallidus and pulvinar in the left hemisphere. Both monkeys also had large activation clusters in the midbrain and pons.

**Table S3.** Effect sizes and 95% Confidence Intervals of Functional ROIs in hippocampus

| | | | Cohen's d | Confidence Interval |
|---|---|---|---|---|
| Monkey P | 40 Hz | lh | 0.180 | [-0.958 1.278] |
| | | rh | -0.440 | [-1.574 0.778] |
| | 10 Hz | lh | 0.433 | [-0.783 1.564] |
| | | rh | 0.581 | [-0.696 1.755] |
| Monkey C | 40 Hz | lh | -0.287 | [-1.392 0.880] |
| | | rh | -0.393 | [-1.516 0.809] |
| | 10 Hz | lh | 0.295 | [-0.874 1.401] |
| | | rh | 0.242 | [-0.912 1.342] |

**Table S4.** Effect sizes and 95% Confidence Intervals of seeds used in seed-based analysis in Figure 6

| **Monkey P** | | 1 | 2 | 3 | 4 | 5 | 6 | 7 | 8 | 9 |
|---|---|---|---|---|---|---|---|---|---|---|
| HP | Cohen's d | -0.189 | -0.296 | 0.078 | -0.669 | -0.746 | -0.320 | 0.211 | 0.185 | -0.319 |
| | CI | [-1.287 0.951] | [-1.403 0.873] | [-1.039 1.177] | [-1.877 0.647] | [-1.987 0.607] | [-1.430 0.857] | [-0.934 1.311] | [-0.954 1.283] | [-1.429 0.857] |
| TE | Cohen's d | -0.585 | 0.431 | 0.409 | 0.146 | 0.336 | 0.497 | -0.738 | 0.340 | -0.178 |
| | CI | [-1.761 0.693] | [-0.785 1.561] | [-0.798 1.535] | [-0.985 1.243] | [-0.846 1.448] | [-0.744 1.645] | [-1.975 0.611] | [-0.843 1.453] | [-1.276 0.959] |

| **Monkey C** | | 1 | 2 | 3 | 4 | 5 | 6 | 7 | 8 | 9 |
|---|---|---|---|---|---|---|---|---|---|---|
| HP | Cohen's d | -0.224 | -0.280 | 0.256 | 0.033 | 0.205 | 0.233 | 0.275 | 0.443 | -0.418 |
| | CI | [-1.323 0.925] | [-1.385 0.884] | [-0.902 1.358] | [1.077 1.135] | [-0.939 1.303] | [-0.919 1.333] | [-0.888 1.379] | [-0.777 1.576] | [-1.545 0.793] |
| TE | Cohen's d | 0.206 | 0.168 | 0.367 | 0.745 | 0.413 | 0.241 | 0.720 | -0.130 | -0.090 |
| | CI | [-0.934 1.305] | [-0.967 1.266] | [-0.825 1.485] | [-0.608 1.985] | [-0.796 1.539] | [-0.620 1.949] | [-1.227 0.997] | [-1.227 0.997] | [-1.189 1.029] |
